# Mechanical Interactions between a Cell and an Extracellular Environment Facilitate Durotactic Cell Migration

**DOI:** 10.1101/460170

**Authors:** Abdel-Rahman Hassan, Thomas Biel, Taeyoon Kim

## Abstract

Cell migration is a fundamental process in biological systems, playing an important role for diverse physiological processes. Cells often exhibit directed migration in a specific direction in response to various types of cues. In particular, cells are able to sense the rigidity of surrounding environments and then migrate towards stiffer regions. To understand this mechanosensitive behavior called durotaxis, several computational models have been developed. However, most of the models made phenomenological assumptions to recapitulate durotactic behaviors, significantly limiting insights provided from these studies. In this study, we developed a computational biomechanical model without any phenomenological assumption to illuminate intrinsic mechanisms of durotactic behaviors of cells migrating on a two-dimensional substrate. The model consists of a simplified cell generating contractile forces and a deformable substrate coarse-grained into an irregular triangulated mesh. Using the model, we demonstrated that durotactic behaviors emerge from purely mechanical interactions between the cell and the underlying substrate. We investigated how durotactic migration is regulated by biophysical properties of the substrate, including elasticity, viscosity, and stiffness profile.

## INTRODUCTION

Cell migration is a fundamental process in biological systems, playing an important role for diverse physiological processes including morphogenesis ^1, 2^ and wound healing ^3^. To illuminate mechanisms of cell migration, various in vitro experimental systems have been employed. It was shown that cells on a substrate migrate in a random fashion characterized as the persistent random walk (PRW) ^4, 5^. PRW allows cells to explore a surrounding space on the substrate in all directions. However, cells often exhibit persistent migration in a specific direction, rather than purely random motion. It was observed that cell migration is guided by a variation in the potential of adhesions to environments (haptotaxis) ^6^, grooved patterns on a substrate at micro and nano length scales (contact guidance) ^7^, and electric fields (galvanotaxis) ^8^.

Cell migration is also guided by the rigidity of surrounding mechanical environments. For example, spatially heterogeneous stiffness of extracellular matrices (ECMs) and tissues is known to direct cell migration in diverse biological processes including wound healing, development, and pathogenesis ^9–15^. In addition, several in vitro experiments showed that the speed and traction force of migrating cells correlate with substrate stiffness ^16–18^. On a substrate with spatially varying stiffness, cells tend to migrate towards the stiffer region of the substrate ^19–23^. This tendency called durotaxis was found to be dependent on the type and composition of the substrate. The durotactic behavior was also observed in multicellular organisms ^24^. In addition, cells exhibit a more sophisticated form of durotaxis. For example, during intravasation, cancer-associated fibroblasts align collagen fibers in an ECM in a specific direction, and then tumor cells can migrate more efficiently via the aligned fibers ^25, 26^ because the ECM becomes much stiffer in direction of fiber alignment.

To understand mechanisms of the durotaxis, several computational models have been developed with different phenomenological assumptions ^27–32^. In most previous models, the mechanics of a substrate was not considered explicitly; cells in the models sense local stiffness imposed on the substrate somehow and change a certain property, depending on the sensed stiffness. For example, it was assumed that the persistency and speed of migration ^31^, cell spreading ^28^, cell traction forces ^30^, or adhesion strength ^32^ vary as a function of local substrate stiffness. Although these models succeeded to recapitulate durotactic behaviors thanks to the phenomenological assumptions, insights provided from these models are inevitably limited because the models cannot explain how cells sense local stiffness and make decisions based on the sensed stiffness. Since mechanical interactions between cells and a substrate are likely to give rise to the durotaxis, the mechanics and deformation of the substrate should be reflected in the model in order to illuminate the intrinsic mechanisms of the durotactic behaviors of cells. Although a few recent models attempted to explicitly account for the mechanical interactions ^29^, the capability and versatility of those models are also limited.

In this study, we developed a computational biomechanical model without any phenomenological assumption to study durotactic behaviors of a cell migrating on a two-dimensional substrate. The model consists of a cell simplified into two points generating contractile forces and a deformable substrate coarse-grained into an irregular triangulated mesh. Using the model, we demonstrated that durotactic behaviors emerge from purely mechanical interactions between the cell and the underlying substrate. We investigated how durotactic migration is regulated by biophysical properties of the substrate, including elasticity, viscosity, and stiffness profile.

## METHODS

We developed a biomechanical model for durotactic cell migration as described in detail below. Table S1 shows the list of all parameters and their values used in simulations.

### Simplification of a cell and a substrate

A migrating cell is simplified into a machine consisting of front and rear cell-points (Fig. 1a). The cell polarity is represented by a vector connecting from the rear cell-point and to the front one. Each cell-point has an adhesion region with a partial donut shape which is defined by outer (*R*_out_) and inner radii (*R*_in_) as well as an angular span (*θ*) defined with respect to the cell orientation. The outer and inner radii of the front cell-point are set to be larger than those of the rear cell-point (i.e. *R*_F,out_>*R*_R,out_ and *R*_F,in_>*R*_R,in_) to reflect asymmetric spreading areas of polarized mesenchymal cells. The angular span of the front (*θ*_F_) and rear adhesion (*θ*_R_) areas is fixed at 180°.

**Figure 1.**
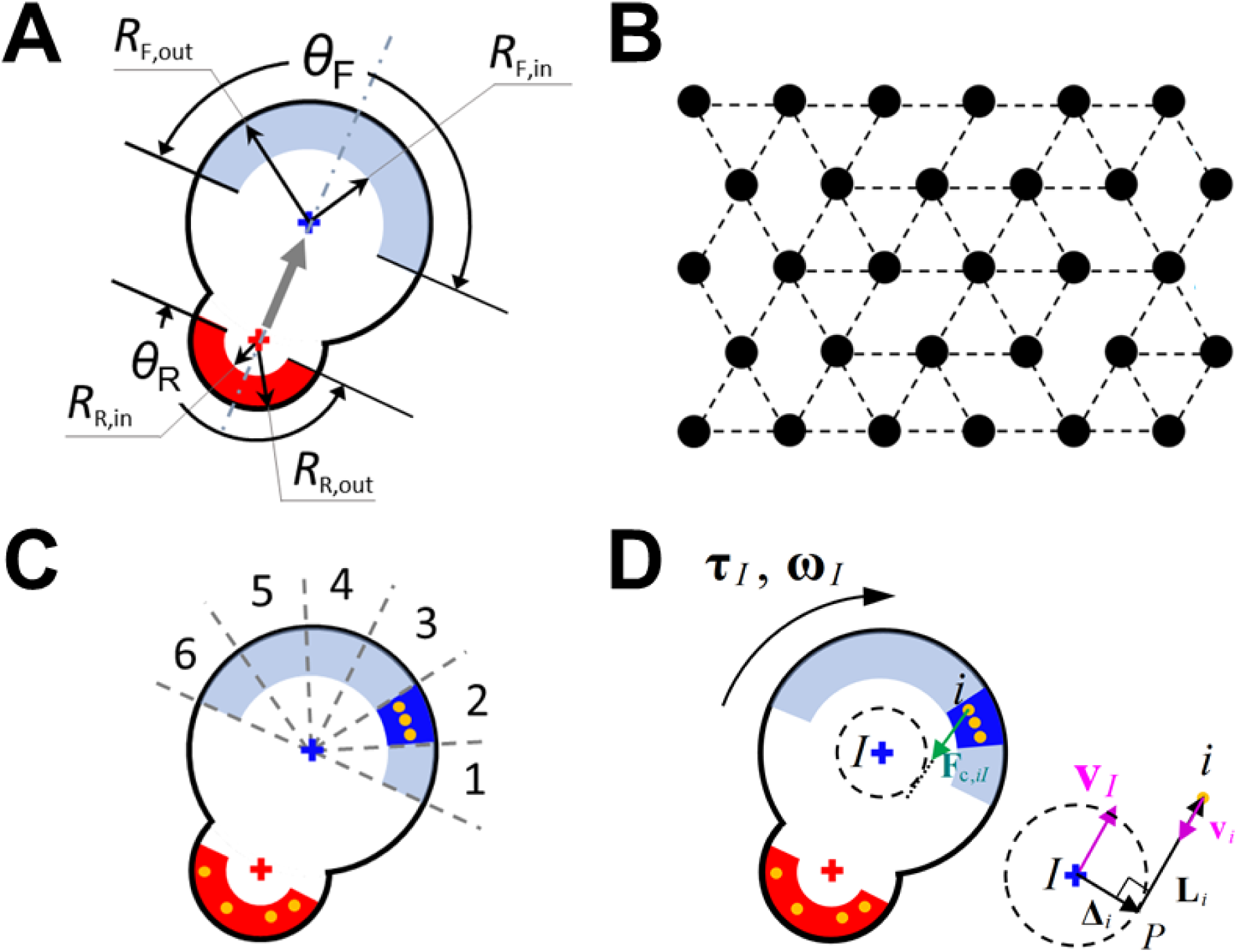
Coarse-grained models for a cell migrating on a substrate. (a) A migrating cell is simplified into a machine consisting of front (F) and rear (R) cell-points. A dot-dashed line represents the direction of cell orientation, and a gray arrow is a vector indicating cell polarity. Each cell-point has an adhesion region with a partial donut shape which is defined by outer (*R*_out_) and inner radii (*R*_in_) as well as an angular span (180°) defined with respect to the orientation. (b) A substrate is modeled as a set of points which are initially located on an equilateral triangular lattice and connected by chains. To account for the polymeric nature of an ECM structure, a fraction of the chains are removed at a probability, *p*. (c) To reflect the dynamic lamellipodial activity, the front adhesion region is partitioned into 6 angular sections by assuming that each of them represents potential lamellipodium. One of the potential lamellipodia is activated randomly and remains active for 6 min. If a lamellipodium reaches its lifetime, it is deactivated, and other potential lamellipodium is activated randomly. By contrast, the entire part of the rear adhesion region remains active all the time. Substrate points that are located within the active adhesion regions immediately become focal adhesion (FA) points (orange). (d) Each cell-point exerts pulling force (**F**_c,*iI*_) on its FAs which originates from constant torque (**τ**_*I*_). As a green arrow indicates, the force vector is directed from the FA point to a tangential point, *P*. **Δ**_*i*_ is a vector connecting the cell-point to *P*, and **L**_*i*_ is a vector connecting *P* with the FA point. The velocity (**v**_*I*_) and angular velocity (**ω**_*I*_) of the cell-point and the velocity of substrate point (**v**_*i*_) are correlated to **Δ**_*i*_ and **L**_*i*_ by the kinematic constraint (Eq. 12).

To take substrate deformation into account, a substrate is modeled as a set of points which are initially located on an equilateral triangular lattice and connected by chains (Fig. 1b). The number of substrate points per unit area (*n*) is varied to simulate an ECM with different fiber densities. To reflect the polymeric nature of an ECM structure, a fraction of the chains are removed at a probability (*p*) ^33^. Thus, the number of chains on each substrate point ranges between 0 and 6. It is assumed that connectivity between substrate points does not vary over time in simulations. We measured mechanical responses of the substrate under various conditions and compared them with measurements in previous literature to verify our substrate model (Supplementary Text and Figs. S6–S9).

### Dynamic formation of lamellipodia

Polarized mesenchymal cells keep forming lamellipodial protrusion in random directions to explore a surrounding space. It was observed that lamellipodia last for ∼10 min before disappearance ^34^. To reflect the dynamic lamellipodial activity, we partition the front adhesion region into 6 angular sections by assuming that each of them represents potential lamellipodium (Fig. 1c). One of the potential lamellipodia is activated randomly and remains active for *T* = 6 min. While a lamellipodium is active, its direction does not change during its lifetime even if the direction of cell polarity changes. If a lamellipodium reaches its lifetime, it is deactivated, and other potential lamellipodium is activated randomly. By contrast, the entire part of the rear adhesion region remains active all the time.

### Interactions between the cell and the substrate

Interactions between cell-points and substrate points are considered as follows. First, substrate points are partitioned into cell-points, depending on their location. If a substrate point is positioned within the outer radius of only one cell-point *I*, it belongs to the region of the cell-point (**R**_*I*_). If a substrate point is located within the outer radius of more than one cell-point, it belongs to the region of the closest cell-point. A fraction of the substrate points in **R**_*I*_ that are located within the active adhesion region immediately become focal adhesion (FA) points where the cell-point *I* exerts a contractile force.

It is assumed that forces exerted by the cell-point *I* originate from constant torque (Fig. 1d). To reflect characteristics of polarized cells, the torque generated by the front cell-point (*τ*_F_) is assumed to be much greater than that generated by the rear cell-point (*τ*_R_). A FA point *i* within **R**_*I*_ experiences a contractile force exerted by the cell-point *I* (**F**_c,*iI*_), and the cell-point also experiences a reaction force (**F**_c,*Ii*_ = -**F**_c,*iI*_). We assume that **F**_c,*Ii*_ is not a centripetal force for the cell-point *I*. The direction of **F**_c,*Ii*_ is consistent with the direction of contractile forces exerted on FA points by myosin motors estimated from the direction of the actin retrograde flow ^34^. In the model, **F**_c,*Ii*_ acts along a tangent line to a circle centered at the cell-point *I* (Fig. 1d). The magnitude of **F**_c,*Ii*_ is not predetermined but calculated at each time step as explained later.

### Formulation of the system

With the assumption of an over-damped system, force balance is considered for a cell-point *I*:

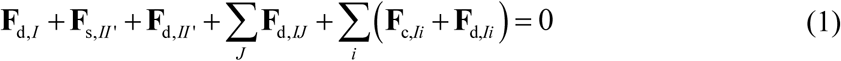

**F**_d,*I*_ is a drag force due to the viscosity of the surrounding medium:

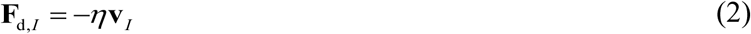

where *η* is a drag coefficient, and **v**_*I*_ represents the velocity vector of cell-point *I*. **F**_s,*II’*_ is the linear spring force derived from a harmonic potential, *U*_s,*II’*_, maintaining the distance between the front and rear cell-points (*r_II’_*) at an equilibrium distance (*r*_0, FR_) via extensional stiffness (*κ*_s,FR_):

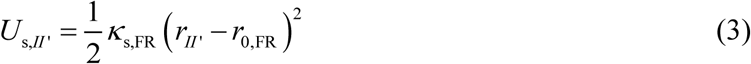

**F**_d,*II’*_ is a viscous drag force between the front and rear cell-points:

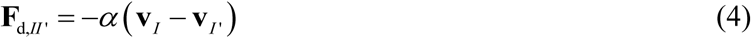

where *α* is a drag coefficient. **F**_d,*IJ*_ is a viscous drag force between a cell-point *I* and other cell-point *J* that belongs to a different cell. The summation for **F**_d,*IJ*_ is calculated for all cell-points located within a critical distance (*r*_crit_) to the cell-point *I*.

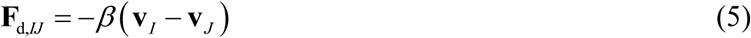

where *β* is a drag coefficient. **F**_d,*Ii*_ is a viscous force acting between a FA point *i* and the cell-point *I*:

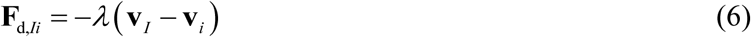

where *λ* is a drag coefficient. The summation for **F**_c,*Ii*_ and **F**_d,*Ii*_ is calculated for all FA points interacting with the cell-point *I*.

In addition, torque balance is considered for each cell-point *I*:

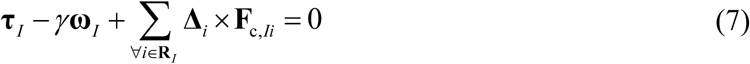

where **τ**_*I*_ is torque generated by the cell-point *I*, *γ* is an angular drag coefficient, ***ω***_*I*_ is the angular velocity of the cell-point *I*, and **Δ**_*i*_ is a vector shown in Fig. 1d.

We consider force balance for a substrate point *i*:

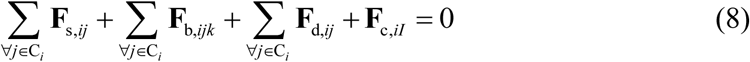

where C_*i*_ indicates the set of all adjacent substrate points connected to the point *i*, and *k* indicates all substrate points connected to either the point *i* or the point *j*. **F**_s,*ij*_ is a linear spring force derived from a harmonic potential, *U*_s,*ij*_, maintaining the length of a chain between the points *i* and *j* (*r_ij_*) at an equilibrium length (*r*_0,M_) via extensional stiffness (*κ*_s,M_):

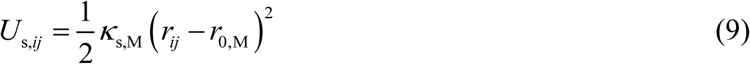

**F**_b,*ijk*_ is a bending force derived from a harmonic potential, *U*_b,*ijk*_, maintaining an angle formed by three points (θ_*ijk*_) at an equilibrium value (θ_*ijk*,0_) via bending stiffness (*κ*_b,M_):

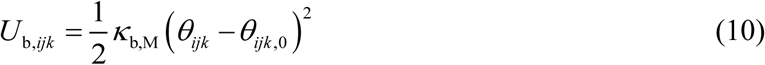

Note that *r*_0,M_ is identical for all chains, whereas θ_*ijk*,0_ can vary depending on the number of removed chains on each substrate point. **F**_d,*ij*_ is a viscous drag force between the substrate points *i* and *j*:

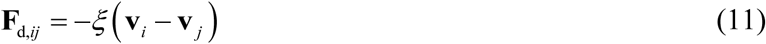

**F**_c,*iI*_ is not zero if a substrate point is a FA point.

As mentioned earlier, the magnitude of the contractile force (**F**_c,*Ii*_) is not predetermined. To calculate |**F**_c,*Ii*_|, we use a kinematic constraint which relates the linear and angular velocities of a cell-point *I* to the velocity of a FA point *i*:

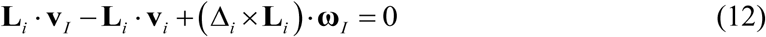

where **L**_*i*_ is shown in Fig. 1d. The kinematic constraint represents that a cell pulls the FA point to propel itself toward that direction, mimicking the mechanism by which cells propel themselves on a substrate in reality. The kinematic constraint corresponds to a situation where the point *i* is pulled by the rotation motion of the circle drawn by a dashed line as if the circle were wrapping around an inextensible string connected to the point *i* (Fig. S1). Collectively, Eqs. 1, 7, 8, 12 are a system of linear equations, so this can be formulated in the form of a matrix equation:

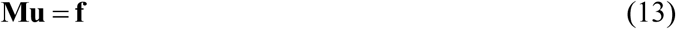

where **M** is a coefficient matrix built based on the relative locations of cell-points and substrate points, **u** is a vector which contains all unknown variables, and **f** is a vector containing constant torques and forces determined by the equations explained above (Fig. S2). The solution of the matrix equation provides the velocities of the cell-points and substrate points and the magnitude of contractile forces acting on FA points at each time step, based on their instantaneous geometrical configurations.

### Variation of substrate stiffness

To study durotaxis, we imposed spatially heterogeneous stiffness on a substrate by varying stiffness in two ways. First, we divide a substrate into two regions and impose different stiffness values on each region by varying extensional stiffness connecting adjacent substrate points (Fig. 2a). We assume that stiffness of the left region is always higher than that of the right region. A cell is initially placed at the interface between two regions with initial orientation toward the +y direction. Second, we assume that the substrate stiffness changes in the x direction with a gradient. The stiffness gradient is discretized into a stepwise decrease in extensional stiffness from the left boundary to the right one (Fig. 2b). When the stiffness gradient is varied between cases, stiffness along the center line at the middle of the substrate in the x direction is fixed at 1 nN/μm. A cell is initially placed on the center line with initial orientation toward either −x or +x direction. Unlike spatially varying extensional stiffness, the removal probability (*p*), the viscosity for drag forces acting between substrate points (*ξ* in Eq. 11), and the areal density of substrate points are spatially uniform. In addition, all parameter values for the cell are kept constant.

**Figure 2.**
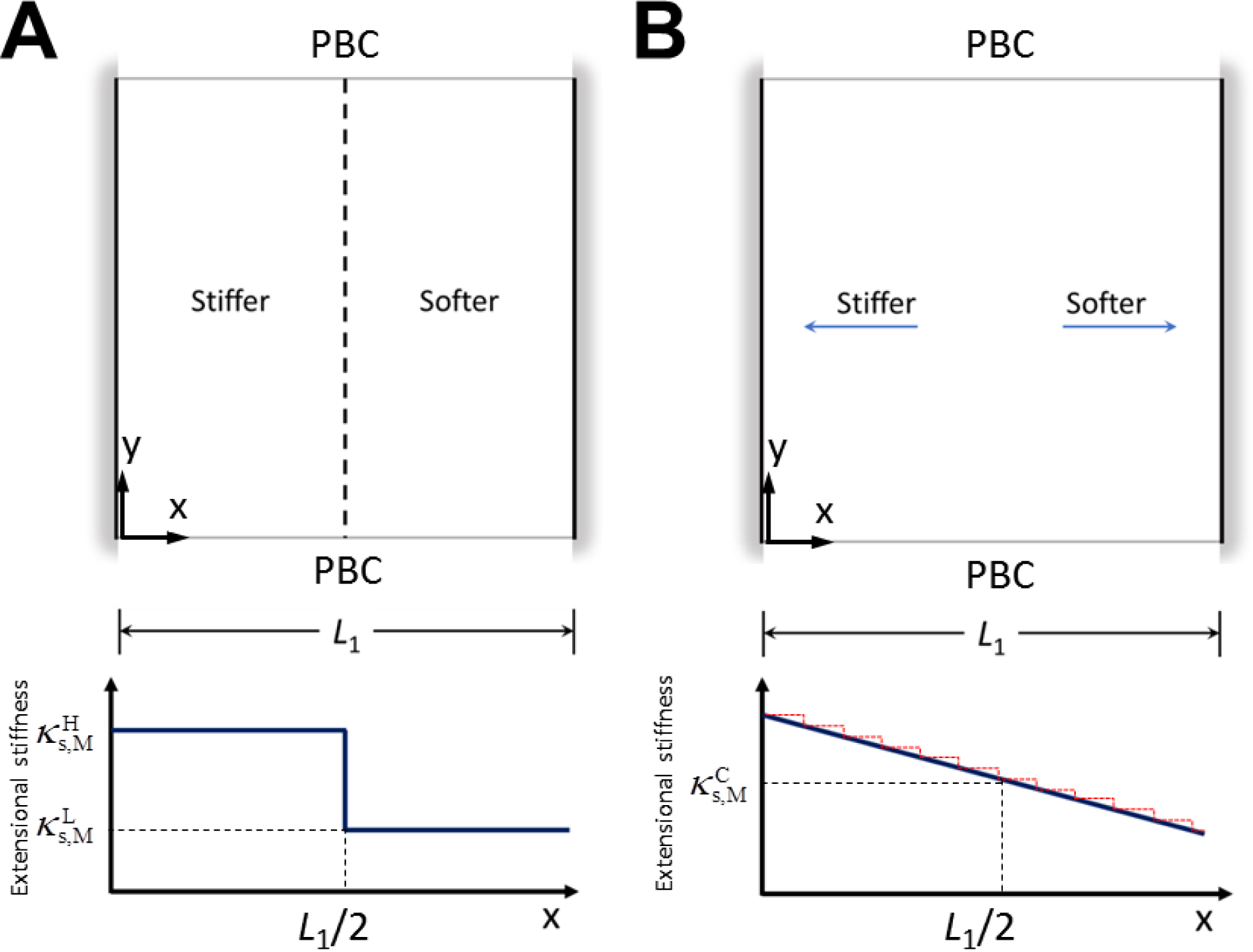
Two types of durotaxis simulations. The width of a square computational domain is *L*_1_, and a periodic boundary condition (PBC) is applied between the top and bottom boundaries of the domain to simulate an infinitely large substrate in the y direction. (a) A substrate with a sharp transition between two different stiffness values (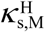 and 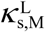) at the center line. (b) A substrate with a stiffness gradient. The gradient (thick solid line) is approximated into a step-wise decrease in the stiffness from the left to the right (red dashed line). The stiffness at the center line 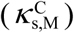 is fixed at 1 nN/μm in all cases.

### Quantification of durotactic migration

To quantify durotactic migration, we calculate the durotactic index (DI). We discretize the trajectory of a migrating cell into multiple short displacements. Each displacement corresponds to the time interval of 0.18 min. DI is defined as the ratio of the number of displacements oriented toward the −x direction to the total number of displacements. DI ∼ 1 indicates strong durotactic migration, whereas DI = 0.5 implies unbiased, random migration.

## RESULTS

### Elasticity and viscosity of a substrate regulate durotaxis

We model a substrate as a material with tunable elasticity and viscosity. Elastic material stores energy well, whereas viscous material dissipates energy. Thus, it is expected that they would play a very different role for the durotaxis. We investigated effects of the elasticity and viscosity of a substrate on durotaxis, using the substrate with two different stiffness values (Fig. 2a). The elasticity is varied by changing the extensional stiffness of the softer region 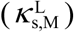 while the ratio of 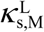 to the stiffness of a stiffer region 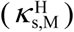 is fixed at 100:1. The viscosity is altered by changing the value of *ξ* in Eq. 11, and *ξ* is assumed to be identical in two regions. We ran simulations with a single cell initially located on the interface between two regions with various values of 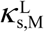 and *ξ* for 12 hr, and then quantified trajectories, DI, and average migration speed. It was found that cells exhibit apparent durotactic behaviors (i.e. migration biased toward the stiffer region) in cases with low *ξ* (Figs. 3a, b). Note that a cell in our model forms lamellipodia in all directions within the front adhesion region at equal probability, and the active lamellipodium lasts during its lifetime. In our previous study, we showed that the cell with dynamic formation of lamellipodia exhibits PRW on a non-deformable substrate ^35^. On a deformable substrate, these cells do not change any of their properties in response to local stiffness. Nevertheless, mechanical interactions between the cell and the heterogeneous mechanical properties of the deformable substrate induce durotactic migration.

**Figure 3.**
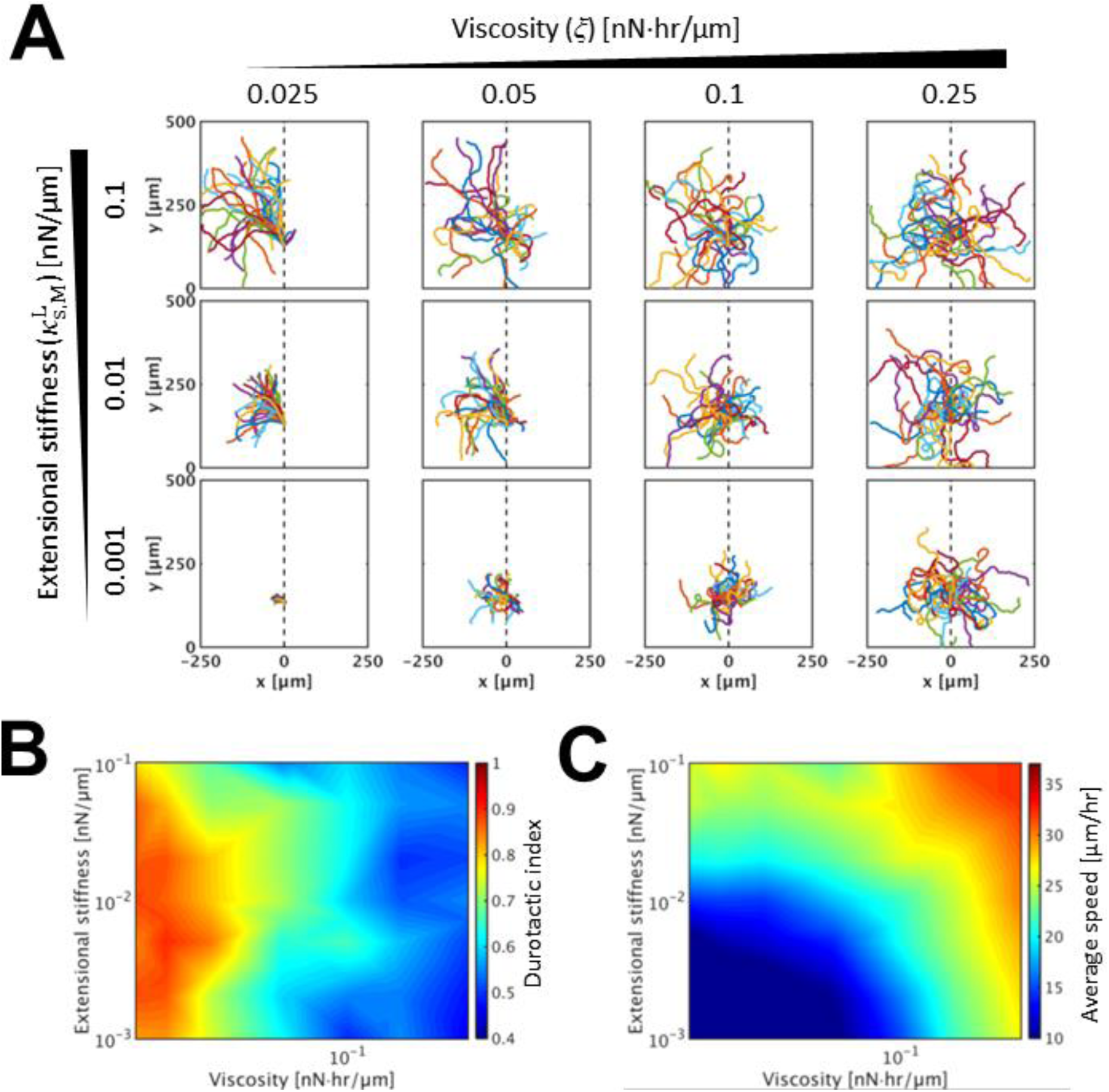
Effects of extensional stiffness and viscosity on durotactic behaviors. In each simulation, a single cell migrates on a substrate with two regions having different stiffness values as shown in Fig. 2a. The stiffness of the left half is always 100-fold higher than that of the right half. We vary the extensional stiffness of the softer region 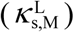 with the ratio of the two stiffness values fixed at 100 (i.e. 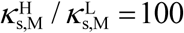). Spatially uniform viscosity, *ξ*, is also varied. Cells are initially located at the interface between two regions with initial orientation toward the +y direction. (a) Trajectories for 12 hr. Each panel contains the trajectories of 30 cells. (b) The durotactic index and (c) average migration speed, depending on extensional stiffness 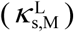 and viscosity (*ξ*).

As *ξ* increases, the bias of trajectories towards the stiffer region and DI are reduced (Figs. 3a, b). With the highest *ξ*, the trajectories are almost unbiased, so DI becomes close to 0.5. This indicates that the enhanced viscosity suppresses durotactic behaviors. The presence of spring and viscous drag forces enables force transmission between interconnected substrate points at short time scales like Kelvin-Voigt material, but the drag force is not sustained at longer time scales. If the viscous effects dominate elastic effects (governed primarily by *κ*_s,M_) in an entire substrate (Fig. S3a, right column), a cell located initially on the interface between two regions cannot distinguish a stiffer region from a softer one because spatially non-uniform elasticity is screened by spatially uniform viscosity. In addition, once a cell enters a softer region, the cell cannot remotely feel the existence of a stiffer region because long-range force transmission is hard to occur in the viscosity-dominated substrate. Thus, it is unlikely for the cell to be reoriented and migrate toward the stiffer region in such a case.

If 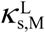 increases from the lowest value to low or intermediate *ξ*, the stiffer region of the substrate can sustain forces for longer time as well as transmit forces better over longer distance because elasticity dominates viscosity (Fig. S3a, left column). By contrast, the softer region is still dominated by viscosity. Then, contractile forces exerted by the cell can be sustained stably only in the stiffer region, so effective durotactic migration can occur from the beginning, leading to the significant enhancement of DI. However, if 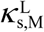 increases further, DI is reduced because the cell can also exert contractile forces on the softer region with elasticity comparable to viscosity, not only on the stiffer region. As a result, DI shows biphasic dependence on 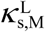 (Fig. 3b).

To confirm our hypothesis, we measured elastic energy stored in two regions in a case with high 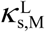 and low *ξ* (Fig. S3b). Energy in the elasticity-dominated region (left half) is much higher and sustained better, whereas energy in the region with elasticity comparable to viscosity (right half) is lower and not sustained well. This contrast between the two regions results in a better environment for a cell to exhibit strong durotactic behaviors. In addition, to directly evaluate the mechanical properties of the substrate, we stretched the substrate without any cell in the −x and +x directions simultaneously to an equal extent (5%) with the center line fixed (Fig. S3c). We used high 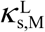 and low *ξ*. As expected, the elasticity-dominated region develops and sustains large forces much better than the viscoelastic region.

Unlike DI, average migration speed is proportional to both 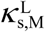 and *ξ* (Fig. 3c). A cell moves by exerting tensile forces on FA points. Resistance of the substrate points to the tensile forces determines a net displacement for the cell; with higher resistance, the cell can move faster. The resistance originates from both viscous and elastic responses of the substrate because such a displacement takes place at short time scales. Enhancement of cell speed with higher elasticity and viscosity of a substrate is consistent with previous experimental observations ^18, 22^.

In sum, a cell exhibits the best durotactic behavior on a substrate with intermediate elasticity and low viscosity. However, the average migration speed is larger with higher elasticity and viscosity. Different dependence of the durotaxis and average speed on the elasticity and viscosity implies that stiffness-dependent migration speed is not responsible for the durotaxis.

### Durotactic behaviors depend on the extent of a variation in stiffness

We also investigated how the extent of a variation in stiffness at the interface between two regions affects durotactic behaviors. We varied 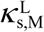 as well as the ratio of two stiffness values 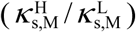 while the viscosity (*ξ*) is kept constant for all simulations. We quantified migration trajectories, DI, and average migration speed. It was observed that the cell migrates faster with larger 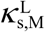 and higher stiffness ratio (Figs. 4a, c). The proportionality of the average speed to 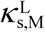 is consistent with results shown in Fig. 3. If the ratio is higher, 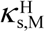 becomes larger. Then, the cell migrates much faster while they migrate on a stiffer region, resulting in higher average speed. DI shows interesting dependence on the stiffness ratio (Fig. 4b). If 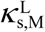 is large, DI reaches a very high value in response to a small increase in the ratio. By contrast, at low 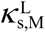, DI slowly increases as the stiffness ratio increases. It was found that DI shows a transition to the high level when 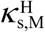 becomes ∼0.15 nN/μm, regardless of 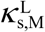. This transition takes place when the elasticity of the stiffer region dominates the viscosity which is constant in all these simulations (Fig. S4). Viscosity still dominates elasticity in the softer region, so the durotactic behaviors become the most efficient. At the lowest 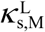, 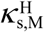 is smaller than the critical value even with the highest ratio, so the high value of DI does not appear. These results confirm that the best durotactic behaviors emerge when a stiffer region is very elastic while the other region is viscous.

**Figure 4.**
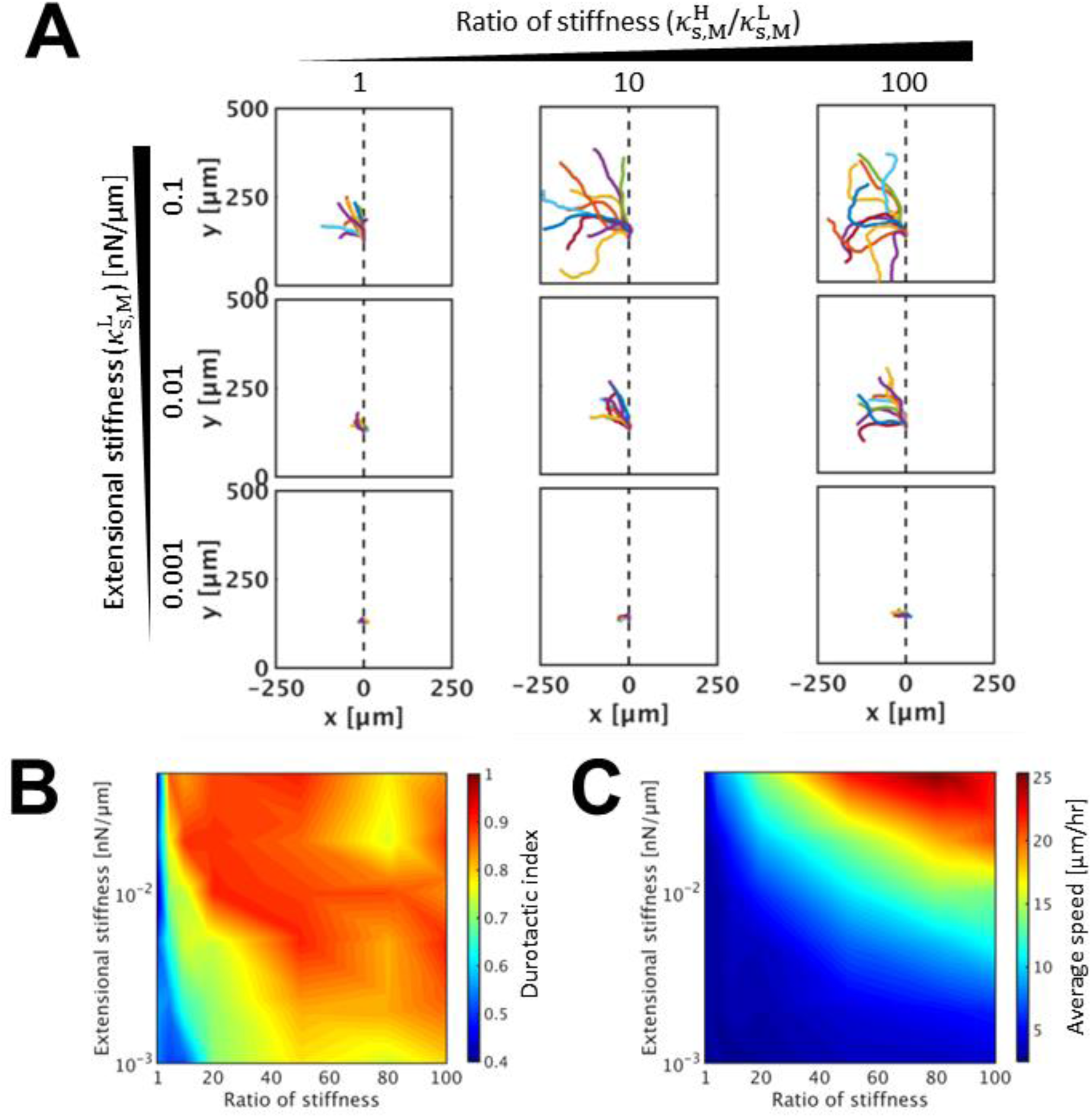
Influences of the extent of a variation in stiffness at the interface. In each simulation, a single cell migrates on a substrate with two regions having different stiffness values as shown in Fig. 2a. We vary the extensional stiffness of the softer region 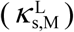 as well as the ratio of two stiffness values 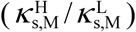. Cells are initially located at the interface between two regions with initial orientation in the +y direction. (a) Trajectories of migrating cells for 12 hr. Each panel contains 10 trajectories. (b) The durotactic index and (c) average migration speed, depending on extensional stiffness 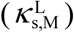 and the stiffness ratio 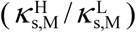.

### Cells can sense a gradual change in the stiffness of a substrate

It was demonstrated above that cells can sense a very sharp transition in substrate stiffness and show durotactic behaviors. We further studied whether or not a cell can also sense a more gradual change in the substrate stiffness. We varied *κ*_s,M_ in a stepwise manner to impose a stiffness gradient (Fig. 2c); *κ*_s,M_ decreases in the +x direction, so chains connecting substrate points initially located in smaller x positions are stiffer than those initially located in larger x positions. The magnitude of the stiffness gradient is changed between 0 and 200 pN/µm^2^, and a cell is positioned on the center of the domain with initial orientation toward either −x or +x direction. Under each condition, we quantified trajectories and the fraction of cells that reach the right boundary. Without any stiffness gradient, most of the cells tend to show PRW and migrate within a region located in the direction of initial cell orientation (Figs. 5a, b). For example, about half of cells initially oriented in the +x direction reached the right boundary, but most of the other cells still migrate within the right half of the substrate (Fig. 5c). In addition, only a small portion of cells initially oriented in the −x direction reached the right boundary. With a small stiffness gradient, overall durotactic behaviors do not change significantly. By contrast, with a high stiffness gradient, cells initially oriented in the +x direction change their migration direction and reach the left boundary at a higher probability. Such trajectories with reorientation are reminiscent of previous experimental results ^19–21^. If the gradient is high enough, none of the cells initially oriented in the +x direction reach the right boundary because all cells are reoriented toward the −x direction. On the other hand, all of the cells initially oriented in the −x direction reach the left boundary because reorientation toward the +x direction is inhibited by the strong durotactic cue. As another measure of the durotactic behaviors, we also quantified how far cells initially oriented in the +x direction travel before the first reorientation toward the −x direction (Fig. 5d). The distance traveled by cells before reorientation is inversely proportional to the stiffness gradient, which is consistent with their trajectories.

**Figure 5.**
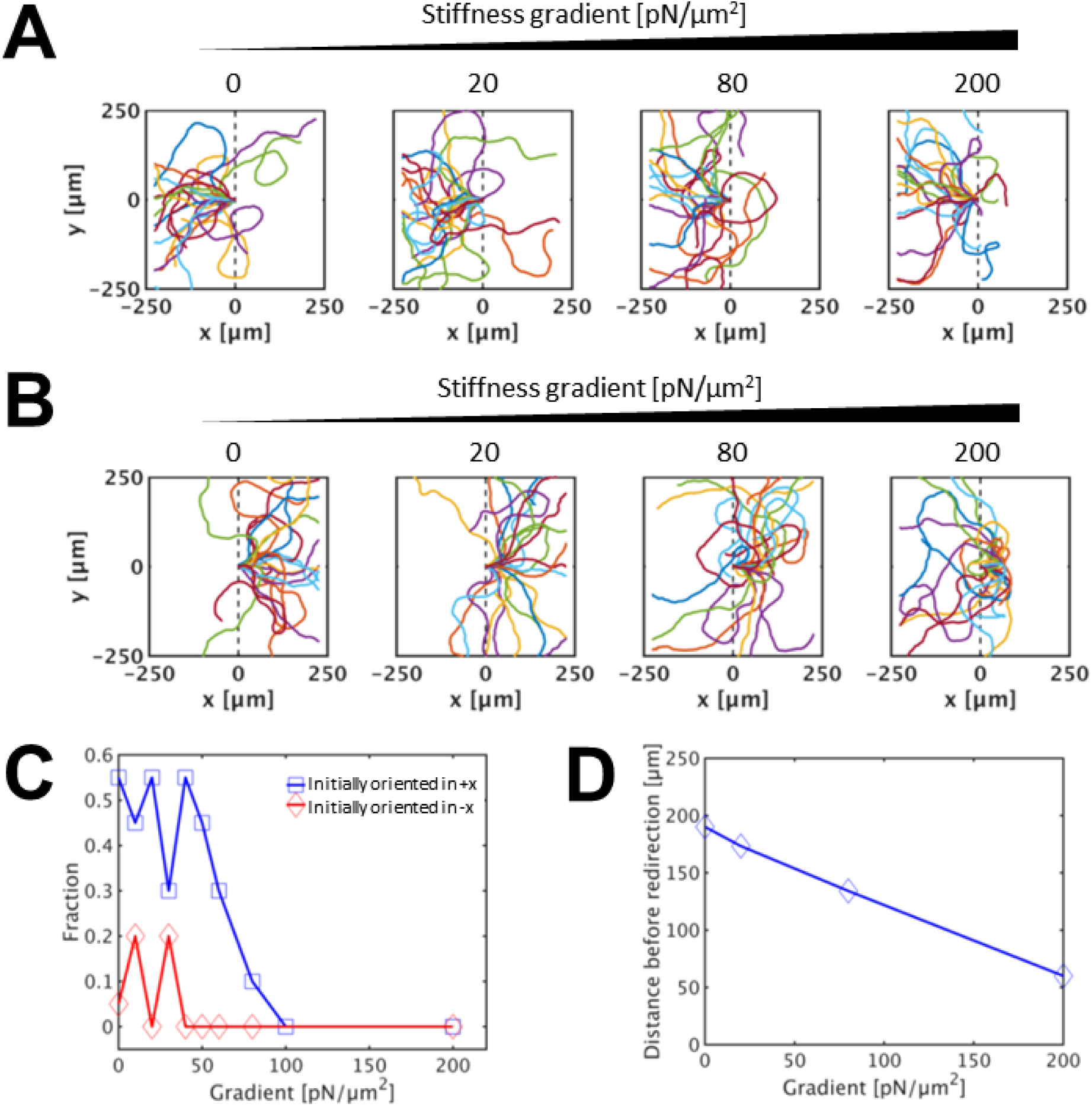
Effects of a stiffness gradient on migration depending on initial cell orientation. We imposed a stiffness gradient on a substrate, so the substrate stiffness linearly decreases from the left to the right as shown Fig. 2b. The values of stress gradients used range between 0 and 200 pN/µm^2^. Cells are initially located at the center. (a, b) Trajectories of cells for 36 hr with initial orientation toward (a) the −x direction and (b) the +x direction. (c) The fraction of cells reaching the right boundary. (d) Distance which cells initially oriented toward the +x direction travel before they are reoriented toward the −x direction.

We also calculated a correlation between instantaneous migration speed and local stiffness. Figure S5 shows the mean and standard deviation of migration speed of cells initially oriented in −x or +x direction with two different stiffness gradients. With a small stiffness gradient (20 pN/μm^2^), there is no obvious correlation between local stiffness and migration speed in both cases with two different initial orientations (Figs. S5A, B). With a high stiffness gradient (200 pN/μm^2^), instantaneous migration speed is always proportional to the local stiffness, regardless of the initial orientation of cells (Figs. S5C, D). Similarity of migration speed at the same local stiffness between cases under different conditions implies that the speed, which is the magnitude of velocity, does not depend on the instantaneous orientation of cells with respect to the direction of the stiffness gradient. Thus, stiffness-dependent speed cannot be the mechanism of durotaxis.

If the stiffness gradient is very high, the substrate viscosity dominates in a region near the right boundary due to a substantial decrease in *κ*_s,M_. Then, durotactic behaviors become very apparent because the existence of both elasticity-dominated and viscosity-dominated regions is an optimal condition for the durotaxis as shown before. To confirm this, we visualized the spatial distribution of forces on a substrate with high stiffness gradient induced by a migrating cell initially oriented in the +x direction (Fig. 6). It was observed that small forces initially accumulate in a small region around the cell. Note that an asymmetric dipole force pattern appears around the cell because the front and rear cell-points exert larger and smaller forces on adjacent FA points, respectively. As the cell migrates for longer time, the magnitude of the forces is increased and propagated. As expected, the forces globally accumulate more in the left region of the substrate, whereas a region near the right boundary shows negligible forces. As the cell is reoriented more in the −x direction, the accumulation and propagation of the forces are more apparent in the left region. This demonstrates that very efficient durotaxis can emerge from very different mechanical properties of the left (elasticity-dominated) and right (viscosity-dominated) regions.

**Figure 6.**
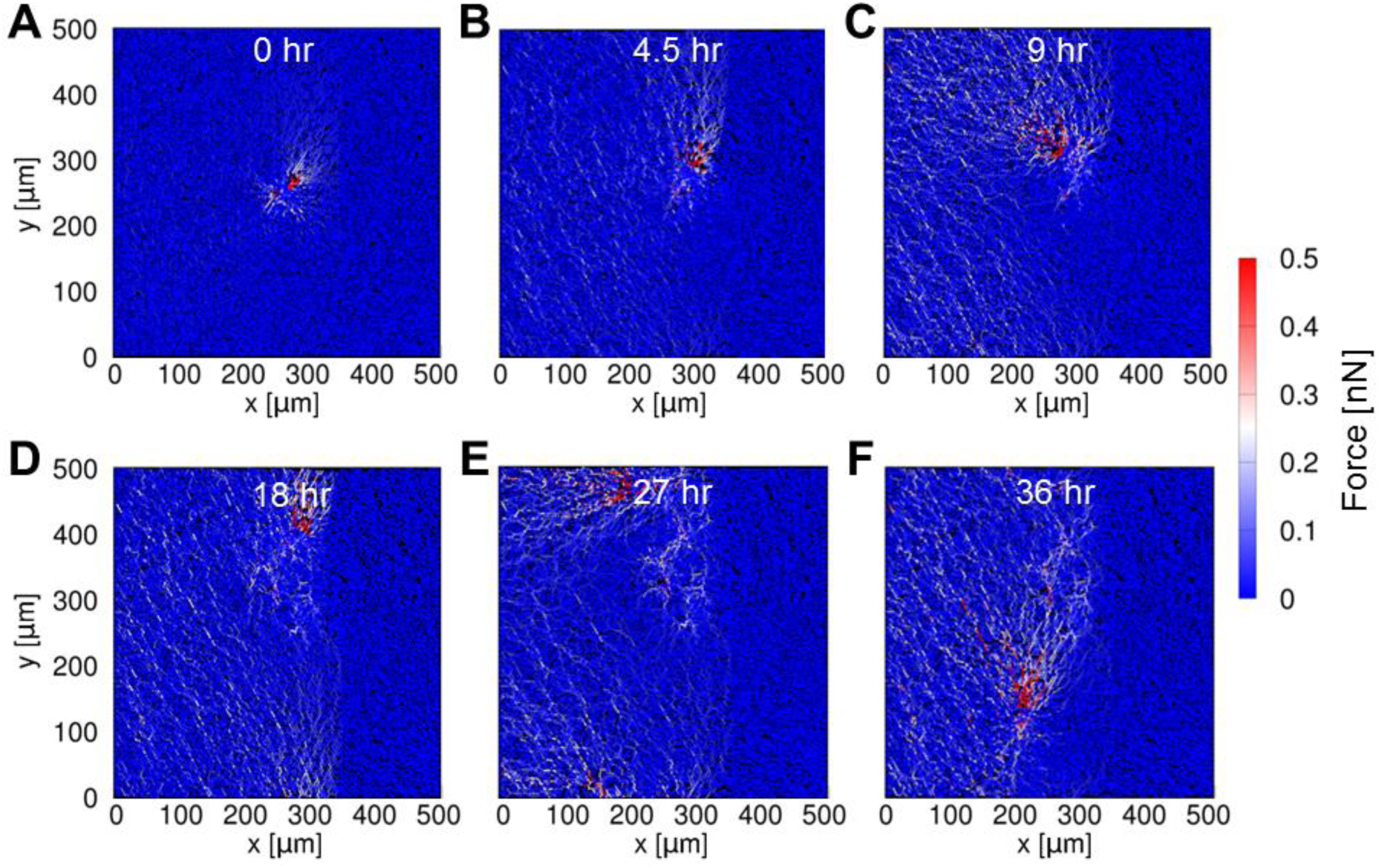
Forces on a substrate with the stiffness gradient of 200 pN/µm^2^ induced by a migrating cell at different time points. Color scaling indicates the amount of forces acting on chains. A cell is located near the center of the red regions with very high forces. At later times, forces accumulate much more in the left region of the substrate, whereas a region near the right boundary shows negligible forces. This asymmetric force development facilitates effective durotactic behaviors.

In sum, we showed that a cell exhibits durotaxis in response to an imposed stiffness gradient. If a durotactic cue induced by a stress gradient is strong enough, the cue can dominate PRW, resulting in the reorientation of cells toward a stiffer region. This cannot be explained by stiffness-dependent migration speed because speed depends only on local stiffness regardless of cell orientation. The durotaxis becomes optimal with high stiffness gradients due to the contrast between mechanical properties of two regions.

## DISCUSSION

Biological and biomimetic materials are often viscoelastic ^36, 37^. Nevertheless, most of the previous experiments studying the durotaxis employed very elastic gels and focused primarily on how a spatial variation in Young’s modulus affects durotactic behaviors. Even in experiments using viscoelastic substrates, relative importance of elasticity and viscosity for durotaxis was not investigated systematically. In this study, we investigated the mechanisms of durotaxis using a model which incorporates a migrating cell with a deformable substrate whose elasticity and viscosity can be controlled separately. We demonstrated that durotactic cell migration toward a stiffer region emerges from purely mechanical interactions between the cell and the substrate.

First, we used a substrate with two regions having different elasticity as in traditional experiments designed for studying durotaxis. We found that the relative importance of the elasticity and viscosity in a substrate determines the efficiency of durotactic migration. If the natures of the mechanical properties of the two regions significantly differ from each other (i.e. elasticity-dominated vs viscosity-dominated), a cell shows a strong tendency to migrate toward the elasticity-dominated region since the elasticity-dominated region can sustain forces exerted by the cell much better than the other.

If elasticity dominates viscosity in both regions, or if one region is elastic with the other region being viscoelastic, the durotaxis still appear, but the efficiency is lower because the cell can exert forces on both regions relatively well. The cell will be displaced to the left significantly when it pulls the stiffer region on the left, but the cell will be displaced back to the right to some extent when it the pulls less stiff region on the right. Thus, the durotactic behaviors are less conspicuous than the case with elasticity-dominated and viscosity-dominated regions. If viscosity dominates in both regions, durotaxis does not emerge because the dominant viscosity of the substrate prevents the cell from sensing the substrate elasticity.

We also imposed stiffness gradually varying with a gradient. It was observed that with high stiffness gradients, all of the cells initially oriented toward a softer region can be reoriented toward a stiffer region. If the gradient is high, viscosity dominates in a region with the lowest stiffness. Thus, the cell shows strong durotactic behaviors because the viscosity-dominated region cannot sustain forces induced by the migrating cell. This explains why the reorientation toward the stiffer region becomes apparent above a critical gradient; at the critical stiffness gradient, the viscosity-dominated region starts appearing near the right boundary of the domain, resulting in effective durotactic behaviors. By contrast, if the stiffness gradient is not high, all regions remain elasticity-dominated, so only weak durotactic behaviors emerge due to local asymmetric distribution of stiffness around the cell.

Our model did not account for complex dynamic behaviors of FA points to keep the complexity of the model minimal; it is assumed that all substrate points that belong to active adhesion regions of the front and rear cell-points immediately become FA points. We showed that the durotaxis can emerge from purely mechanical interactions between a cell and a substrate without force-dependent FA dynamics. However, the importance of FAs for mechanosensitive and durotactic behaviors of cells has been demonstrated in previous studies ^38–43^. Note that our results do not contradict the role of FA dynamics for the durotaxis. Indeed, we plan to incorporate FA dynamics with our model to investigate the role of FAs for durotactic behaviors of migrating cells. It is expected that a cell with FA dynamics in the model will be able to distinguish lower stiffness from higher stiffness better even when all regions of the substrate are elasticity-dominated, leading to enhanced durotaxis.

## CONCLUSIONS

We showed that tuning the mechanical properties of the substrate can suppress or enhance the durotactic behaviors of cells, using a computational model without any phenomenological assumption. Durotaxis is not possible merely through decisions made by cells. Even if a cell makes decisions for migration based on mechanical cues, actions taken by the cell must be executed within the limits of the laws of mechanics. Therefore, only with proper consideration of substrate mechanics, durotaxis can be studied as an emergent behavior caused by the interplay between cell activities and the substrate as we demonstrated.

## ACKNOWLEDGMENTS

The authors gratefully acknowledge the support from the National Institute of Health (1R01GM126256).

## SUPPORTING INFORMATION

### Measurement of the mechanical response of a substrate

Previous studies have shown how the microscopic properties of an extracellular matrix regulate the mechanical properties of the matrix at macroscopic level ^44, 45^. We employed one of the previous computational approaches for the coarse-grained model of a substrate ^45^, so we evaluated the response of the substrate under various conditions to confirm that our substrate model shows behaviors consistent with the previous study. We measured the elastic response of a substrate without cells by applying shear strain to one boundary whose normal direction is +x while the other boundary fixed. Then, we measured stress exerted on the moving boundary as a measure of the mechanical response of the substrate. The stress is calculated by summing forces acting on all substrate points clamped to the moving boundary and dividing the sum by the area of the boundary. To calculate the boundary area in our two-dimensional model, we assume a finite thickness for the substrate, *d* = 1 μm. Thus, the area of the moving boundary is equal to the length of the boundary times the thickness. To understand how each microscopic factor affects the mechanical response of the substrate, we changed each of the four parameters: bending stiffness (*κ*_b,M_), extensional stiffness (*κ*_s,M_), removal probability of chains (*p*), and areal density of substrate points (*n*).

Figure S6 shows the stress and differential modulus of the substrate with various values of *κ*_b,M_. Overall, stress level is larger with higher *κ*_b,M_. With relatively large values of *κ*_b,M_ (≥ 1 nN·μm), stress is directly proportional to strain at low strains, so differential modulus is almost independent of strain. With the smallest bending stiffness, stress is no longer directly proportional to strain. These imply that bending deformation is mainly responsible for the linear response of the substrate at low shear strains. At higher strains, stress diverges, showing strain-stiffening with a non-linear response. Stress-strain curves in all cases converge at high strains, meaning that substrate stiffness becomes similar at high strains. Due to the convergence of curves, stiffening of the substrate at high strains is less conspicuous in cases with larger *κ*_b,M_.

Figure S7 shows how the stress-strain response of the substrate changes depending on the value of *κ*_s,M_. Magnitudes of stress and differential modulus at high strains significantly differ from each other, and stress does not converge but diverge at high strains. This indicates that the responses at high strains originate mostly from the extensional deformations of chains.

Figure S8 shows the stress and differential modulus of the substrate with different values of *p*. Stress and differential modulus tend to be larger with lower *p* since the substrate points are interconnected better. At higher strains, strain-stiffening emerges in all cases, but it is more conspicuous in a case with larger *p*. Stress curves do not converge well at high strains because the substrate connectivity also affects substrate stiffness at high strains.

We further investigated influences of the areal density of substrate points (*n*). Figure S9 shows the stress and differential modulus of the substrate with different values of *n*. It was observed that stress and substrate stiffness are independent of *n*. Thus, with higher *n*, it is possible to simulate a substrate with more FA points and the same stiffness, but computational cost would increase.

In sum, we found that stress exerted on a substrate varies linearly at low strains and shows non-linear stiffening at higher strains. The bending and extensional stiffnesses determine stress and substrate stiffness at low and high strains, respectively. These observations are consistent with previous experimental and computational results ^45^.

## SUPPLEMENTARY FIGURES

**Figure S1.**
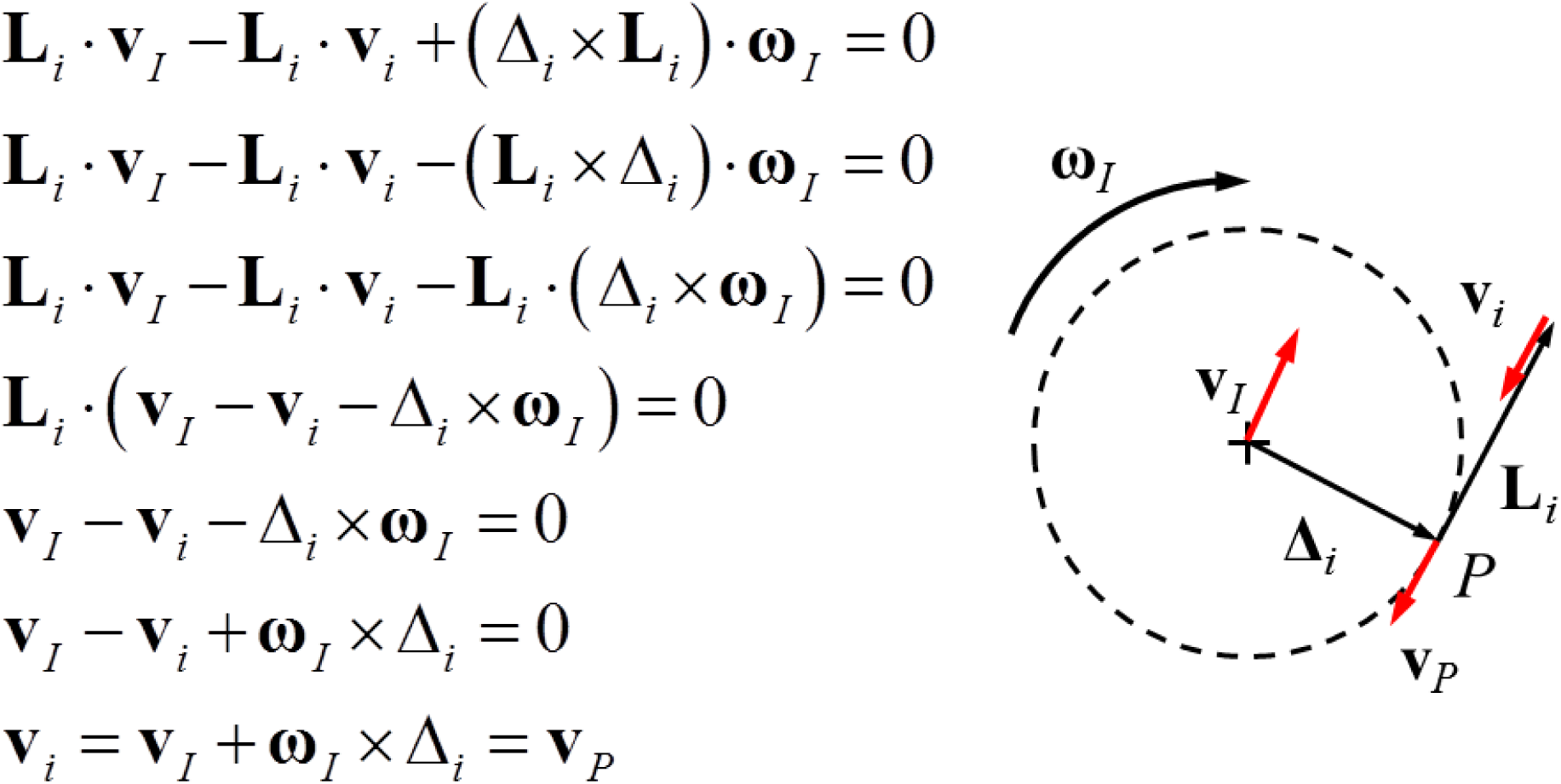
Explanation of the kinematic constraint. For each cell-point, one of the FA points is randomly selected as a reference point (*i*) for the kinematic constraint. It is assumed that the point *i* is pulled by the rotation motion of the circle drawn by a dashed line as if the circle were wrapping around an inextensible string connected to the point *i*. Then, **v**_*i*_ is equal to **v**_*P*_ because the points *i* and *P* are connected by the string. Eq. 12 representing the kinematic constraint essentially means that **v**_*i*_ is equal to **v**_*I*_ + **ω**_*I*_ × Δ_*i*_ which is the same as **v**_*P*_.

**Figure S2.**
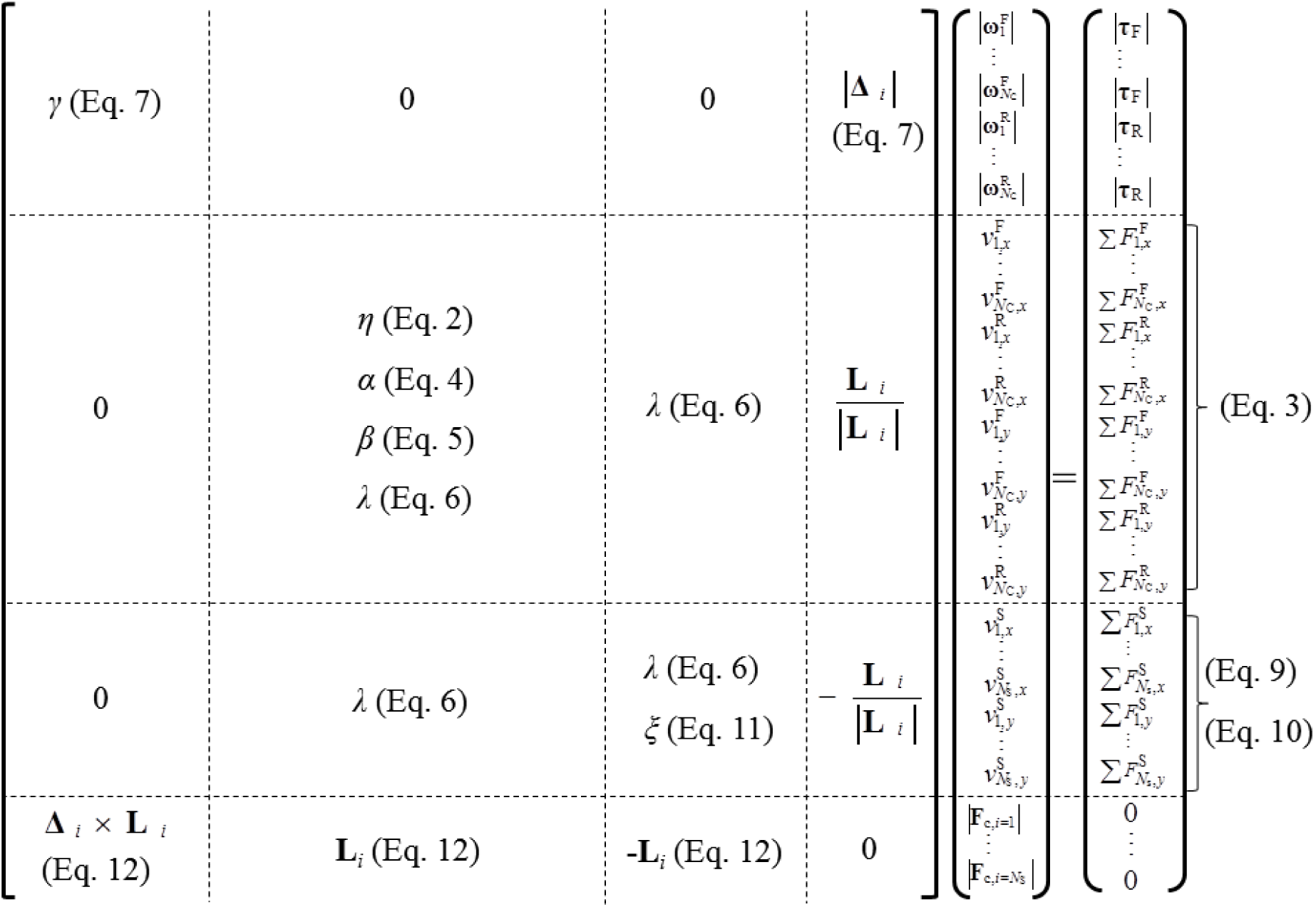
Constitution of the matrix equation. This figure shows elements in the matrix equation (Eq. 13), **Mu** = **f**. Equation numbers are written, wherever possible, to indicate where elements originate from. The vector **u** has the magnitude of **ω**_*I*_ and the x and y components of **v**_*I*_ and **v**_*i*_ as well as the magnitude of contractile forces acting on FA points. *N*_C_ and *N*_S_ are the total number of cell-points and substrate points, respectively. Subscripts F and R indicate the front and rear cell-points, respectively. The vectors **Δ**_*i*_ and **L**_*i*_ are shown in Fig. 1d.

**Figure S3.**
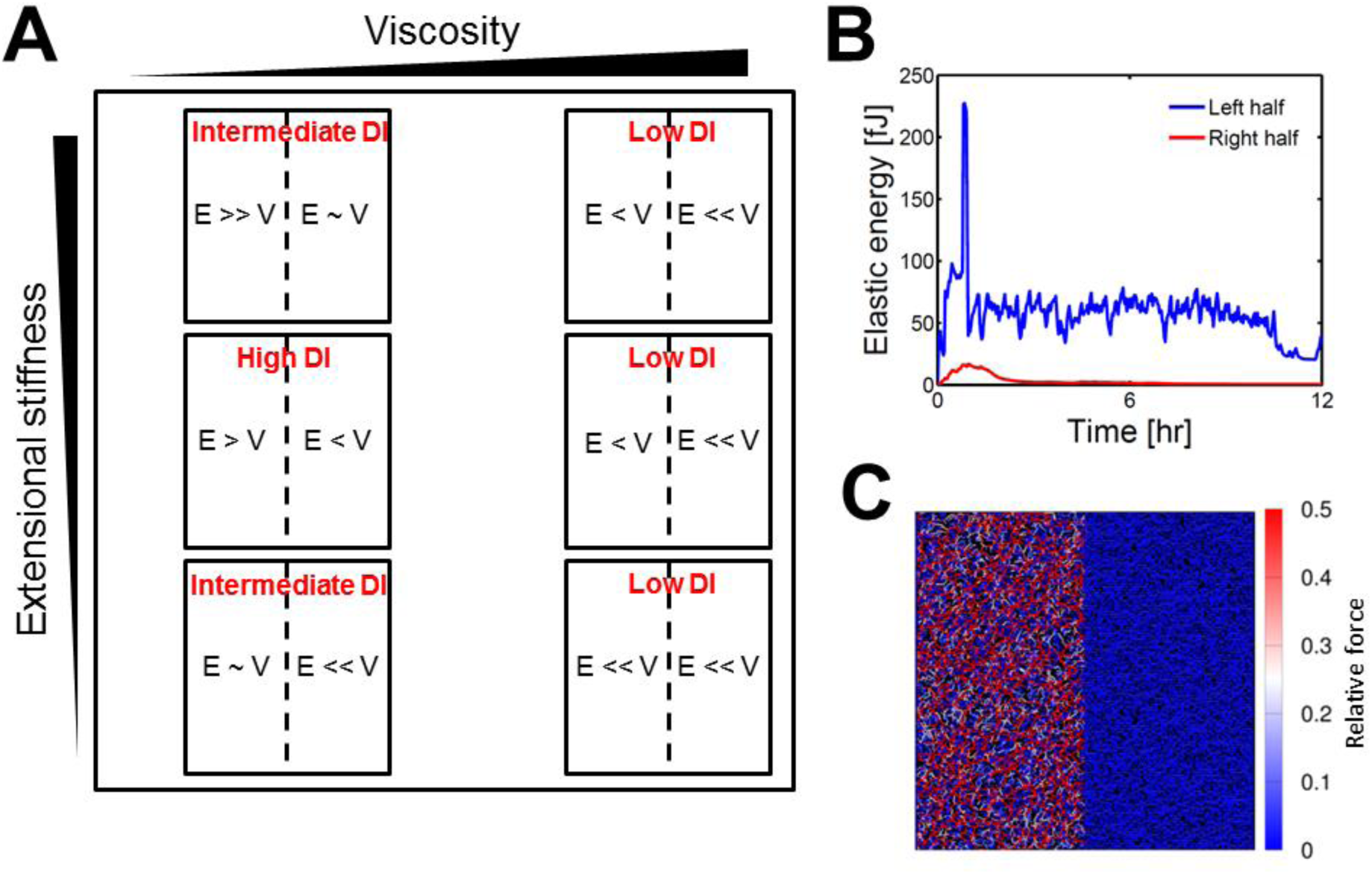
Prediction and quantification of durotactic behaviors. (a) The durotactic index (DI) depending on the relative importance of the elasticity (E) and viscosity (V) in two regions of a substrate. As in Fig. 3, it is assumed that extensional stiffness of the left region is 100-fold higher than that of the right region, whereas substrate viscosity is identical. This explains observations shown in Fig. 3. (b) Elastic energy stored in the two regions in the top-left case shown in Fig. 3A. (c) A snapshot showing the spatial distribution of forces induced by 5% stretch of the substrate in both −x and +x directions to an equal extent. The substrate is initially a square whose side is 0.5 mm. We used the same conditions as those used for the top-left case shown in Fig. 3A. The left half of the substrate is an elasticity-dominated region (E >> V). For (b, c), 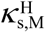 = 10 nN/μm, 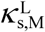 = 0.1 nN/μm, *κ*_b,M_ = 1 nN·μm, *p* = 0.4, and *n* = 4 × 10^4^ mm^−2^.

**Figure S4.**
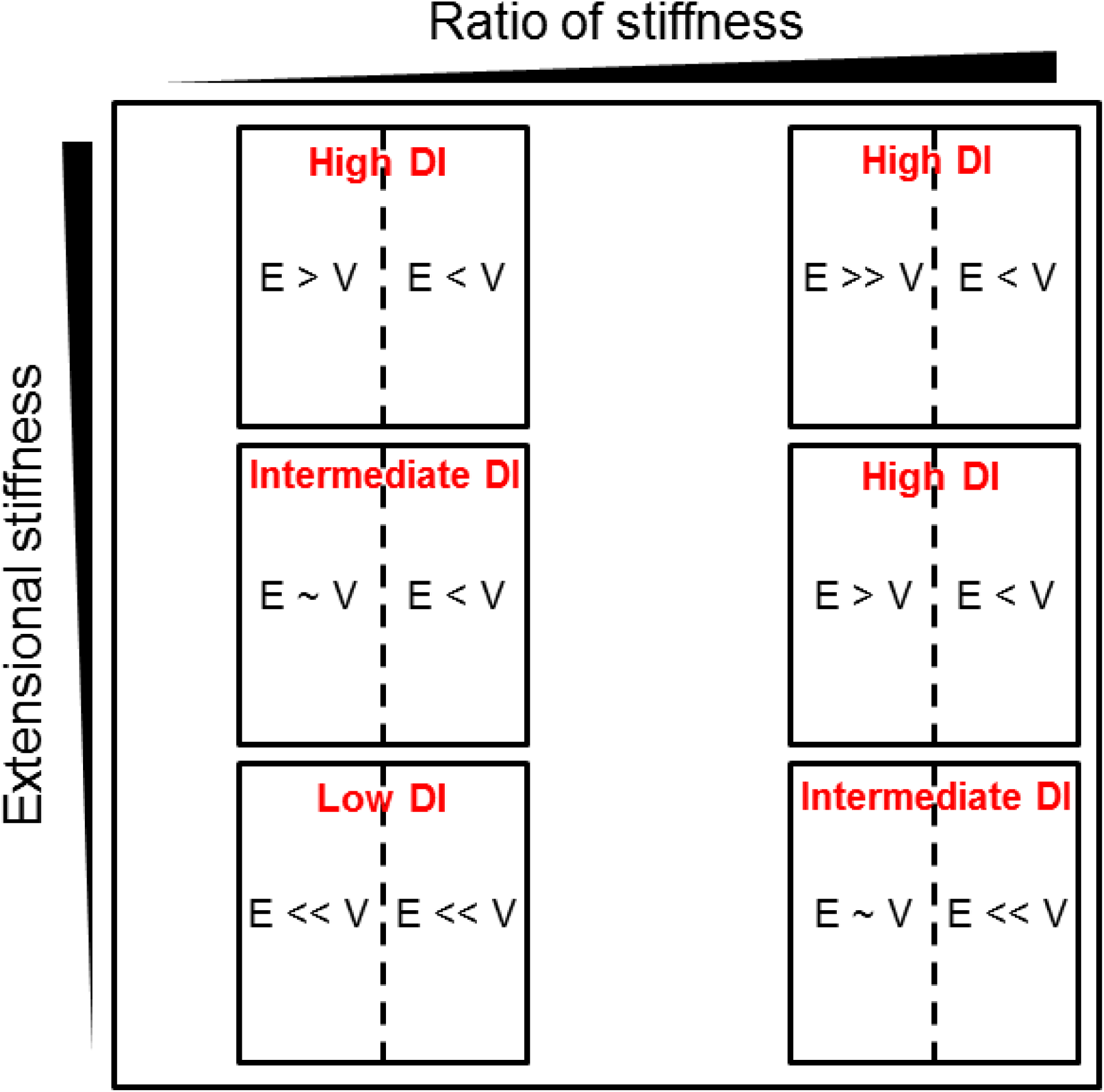
The durotactic index (DI) and the relative importance of the elasticity (E) and viscosity (V) in two regions of a substrate, depending on the extensional stiffness of a softer region 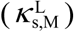 and the ratio of two extensional stiffness values. This explains observations shown in Fig. 4.

**Figure S5.**
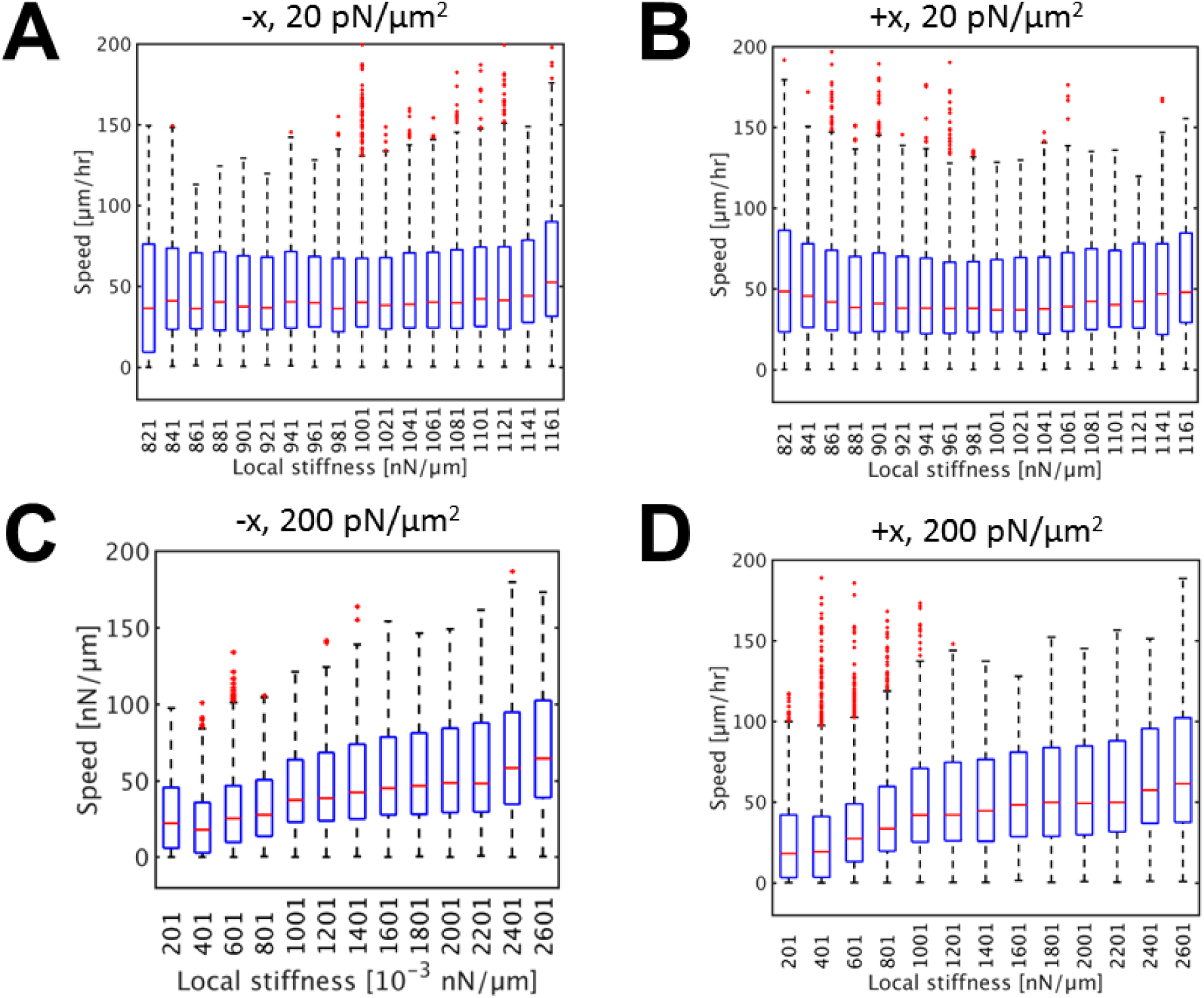
A correlation between instantaneous cell speed and local substrate stiffness. For this quantification, we used cases shown in Fig. 5 with cells initially oriented toward (a, c) the −x direction and (b, d) the +x direction with two different stiffness gradients: (a, b) 20 pN/μm^2^ and (c, d) 200 pN/μm^2^.

**Figure S6.**
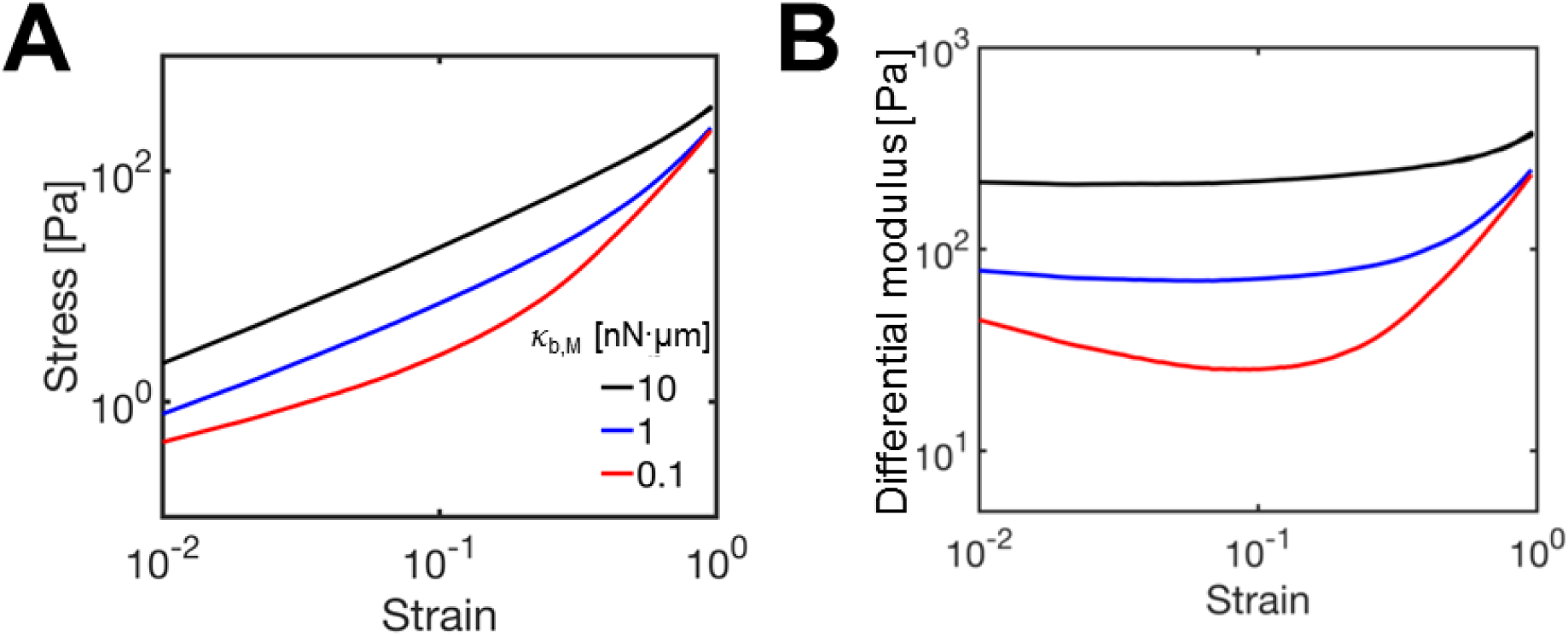
The response of a substrate to increasing shear strain with three values of bending stiffness of the substrate (*κ*_b,M_). (a) Stress and (b) differential modulus of the substrate depending on strain level. *κ*_s,M_ = 1 nN/μm, *p* = 0.4, and *n* = 2 × 10^4^ mm^−2^.

**Figure S7.**
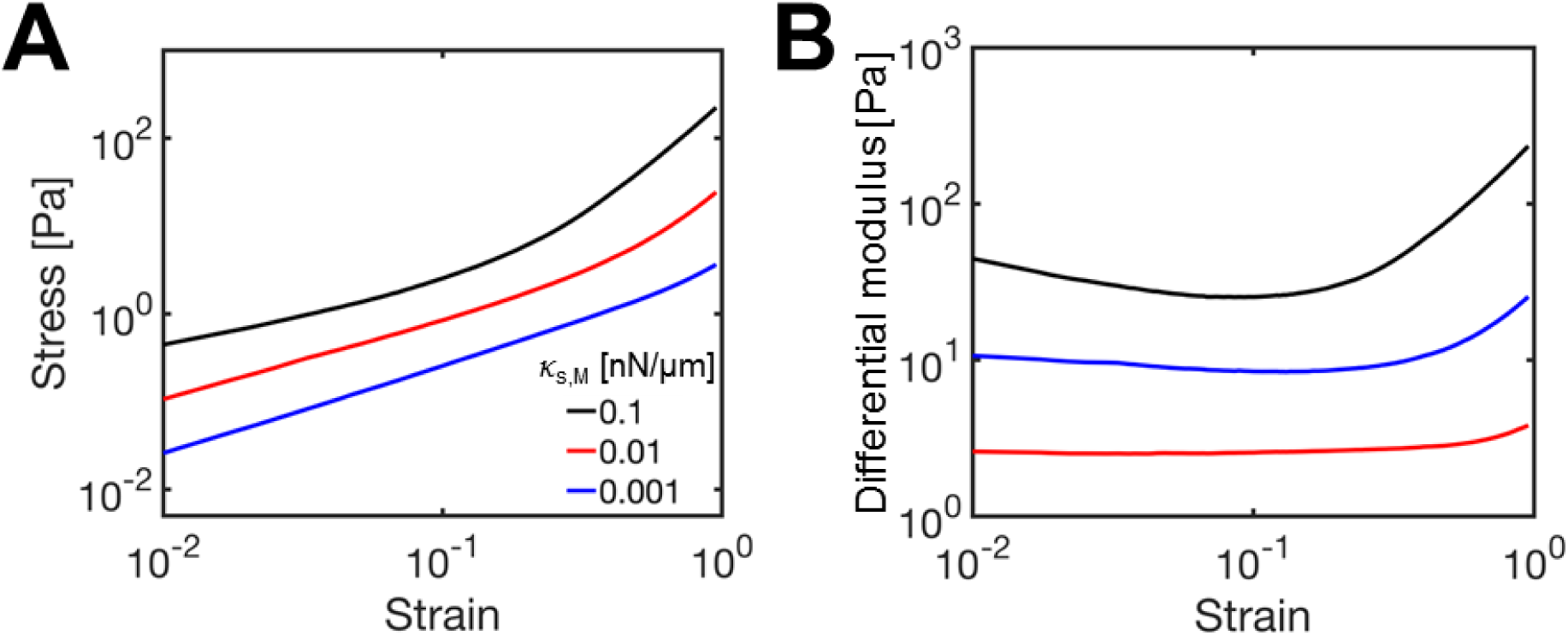
The response of a substrate to increasing shear strain with three different values of extensional stiffness of the substrate (*κ*_s,M_). (a) Stress and (b) differential modulus of the substrate depending on strain level. *κ*_b,M_ = 0.1 nN·μm, *p* = 0.4, and *n* = 2 × 10^4^ mm^−2^.

**Figure S8.**
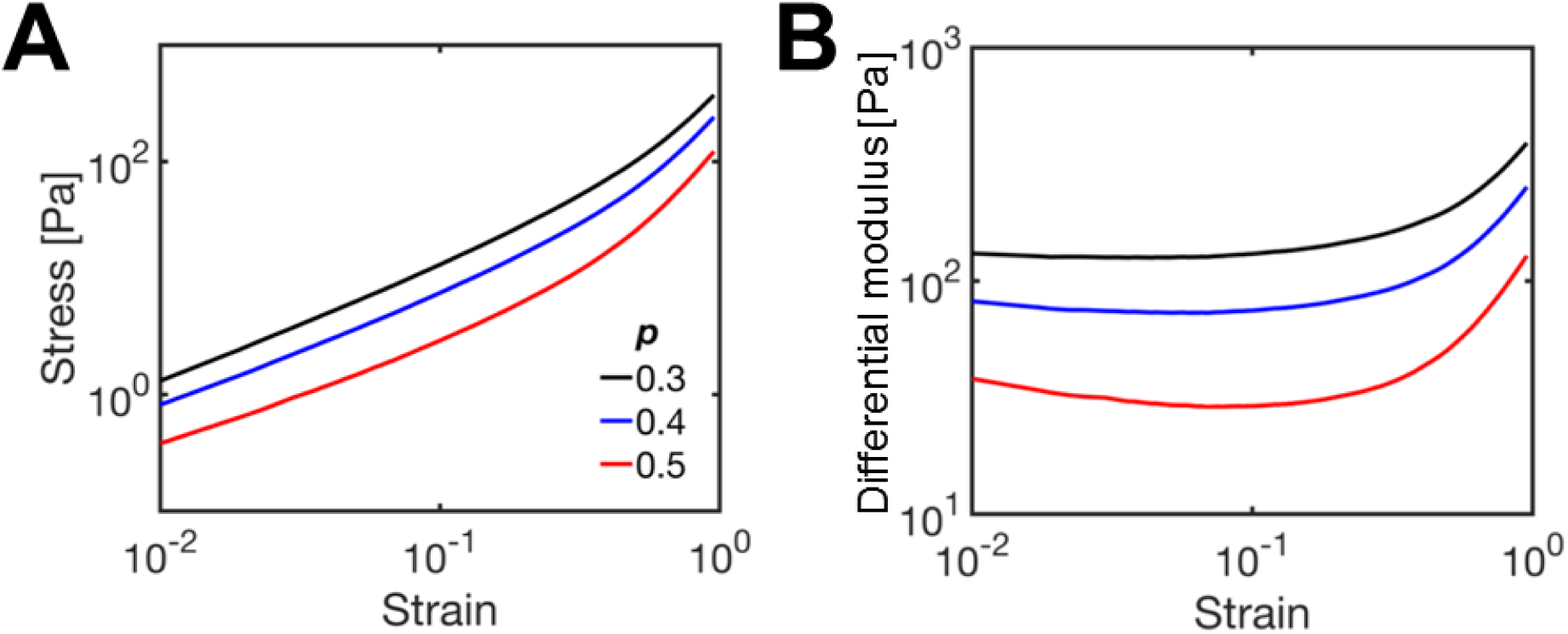
The response of a substrate to increasing shear strain with three different removal probabilities of chains on the substrate (*p*). (a) Stress and (b) differential modulus of the substrate depending on strain level. *κ*_b,M_ = 1 nN·μm, *κ*_s,M_ = 1 nN/μm, and *n* = 4 × 10^4^ mm^−2^.

**Figure S9.**
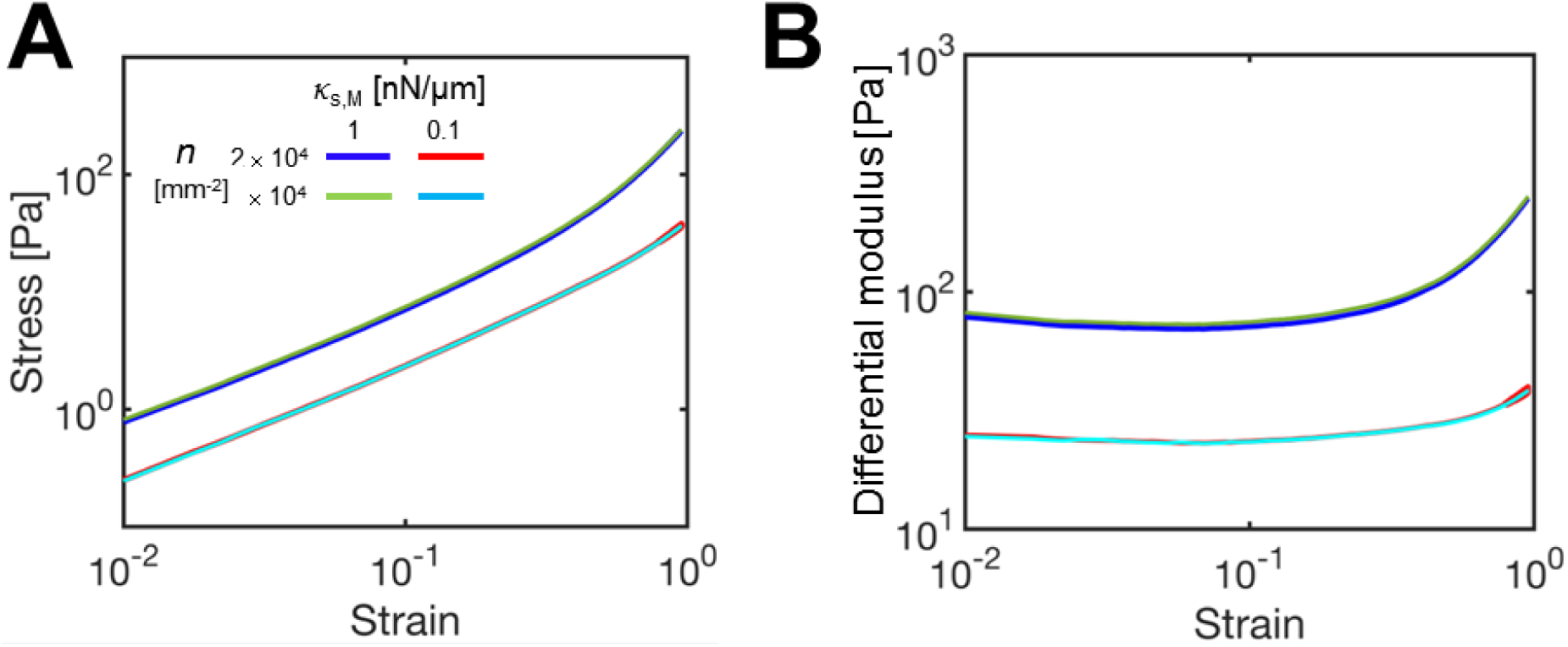
The response of a substrate to increasing shear strain with two different areal densities of substrate points (*n*) and two values of extensional stiffness of the substrate (*κ*_s,M_). (a) Stress and (b) differential modulus of the substrate depending on strain level. *κ*_b,M_ = 1 nN·μm, and *p* = 0.4.

## SUPPLEMENTARY TABLE

**Table S1.**
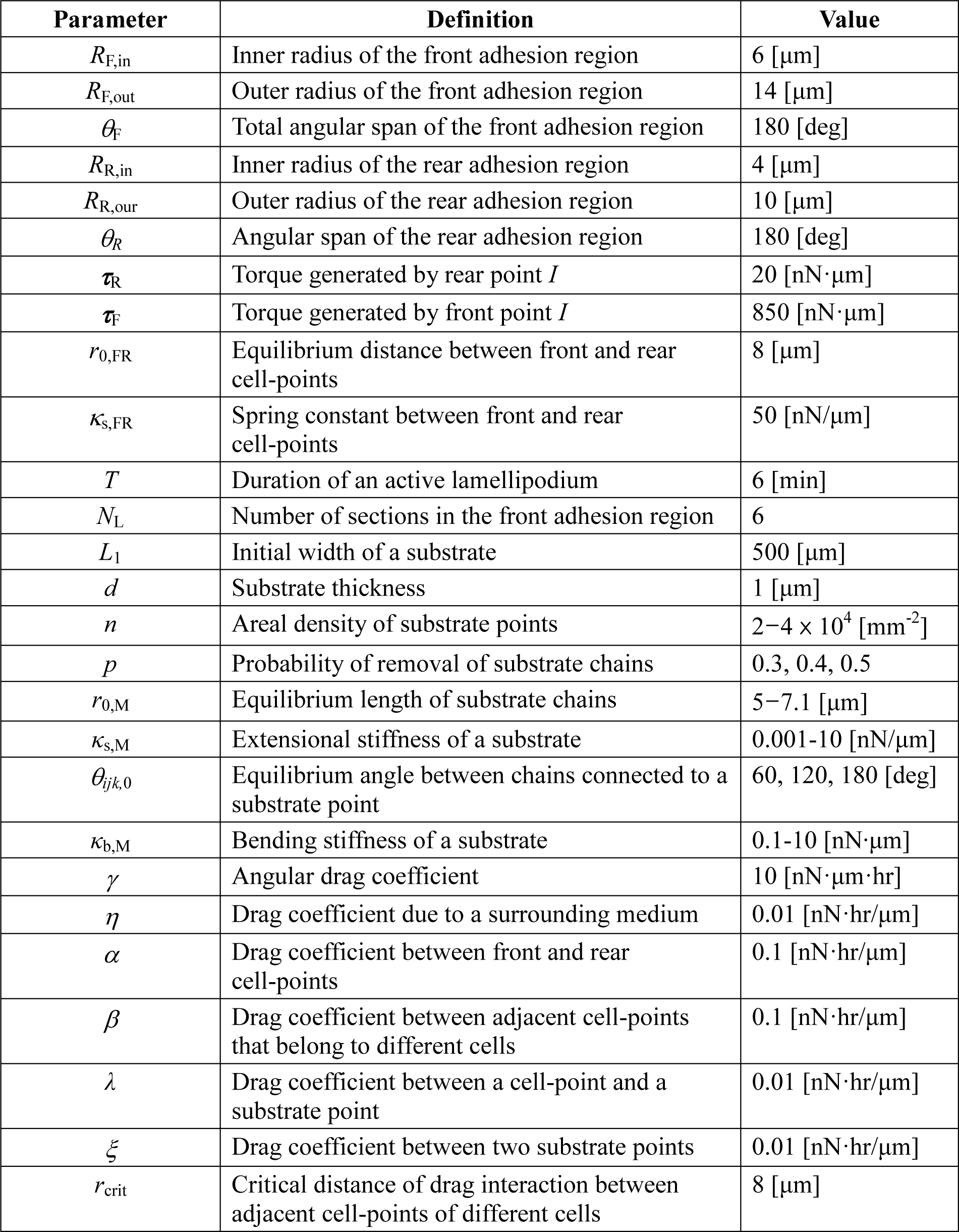
Parameters used in the model and their reference values.

